# BIOMECHANICAL CHARACTERIZATION OF PORCINE LOWER LIMB ARTERIES FOR PRECLINICAL EVALUATION OF PERIPHERAL VASCULAR DEVICES

**DOI:** 10.64898/2025.12.01.691735

**Authors:** Barbara Batista de Oliveira, Frazer Heinis, Anastasia Desyatova, Jason MacTaggart, Alexey Kamenskiy

**Affiliations:** Department of Surgery, University of Nebraska Medical Center, Omaha, NE, USA; Department of Biomechanics, University of Nebraska Omaha, Omaha, NE, USA

**Keywords:** swine, lower extremity arteries, peripheral artery disease, biomechanics, limb flexion

## Abstract

**Purpose:** Peripheral artery disease (PAD) predominantly affects the lower extremities, where complex biomechanical deformations during limb flexion contribute to disease progression and treatment failure. While human and cadaver studies have characterized these deformations, preclinical device testing requires large-animal models that replicate human arterial anatomy and biomechanics. Swine are commonly used, yet their biomechanical comparability to humans remains poorly defined.

**Methods:** We performed a detailed morphometric and biomechanical analysis of the external iliac (EIA), superficial femoral (SFA), and popliteal (PA) arteries in 20 Yucatan and 16 domestic swine using computed tomography angiography. Arteries were evaluated in straight and flexed limb postures to assess diameters, lengths, axial compression, tortuosity, bending angles, and inscribed sphere radii. Breed-specific effects of age and weight were also analyzed.

**Results:** Porcine arterial dimensions closely matched human lower extremity vessels. EIA diameters (4.9-7.2 mm) corresponded to human SFA, porcine SFA (4.1-5.9 mm) approximated human PA, and porcine PA (3.0-4.7 mm) resembled human tibial arteries. Segment lengths supported use of multiple devices. Flexion induced 12-33% axial compression, mimicking worst-case human scenarios. Tortuosity increased distally, and bending characteristics in porcine PAs aligned with human data. In Yucatan swine, vessel diameters were stable with age and weight, while domestic swine exhibited greater variability. Flexion-induced compression and tortuosity were not influenced by age or weight.

**Conclusion:** Swine are well-suited for modeling the geometry and biomechanics of human lower extremity arteries. Their anatomical compatibility and ability to replicate physiologic deformations make them valuable models for preclinical testing of PAD therapies and vascular devices.

## INTRODUCTION

Peripheral artery disease (PAD) is a progressive atherosclerotic condition affecting the arteries of the lower extremities. It is a major contributor to cardiovascular morbidity and mortality worldwide, affecting over 200 million people globally^6^. PAD is particularly prevalent in aging populations and individuals with diabetes, hypertension, or smoking history^26^. It is associated with significant complications, including intermittent claudication, critical limb ischemia, limb loss, and impaired quality of life^6^. Despite advances in medical and surgical therapies, long-term outcomes following endovascular or open revascularization remain suboptimal, with persistently high rates of restenosis, stent and graft failure, and limb amputation^7,20,21,24,30^.

A major challenge in managing PAD stems from the unique biomechanical environment of the lower extremity arteries. These vessels are exposed to complex, dynamic deformations during routine activities such as walking, sitting, and kneeling. Limb flexion in particular induces axial compression, bending, torsion, and changes in arterial curvature^1,17,22,4,19,5^. These mechanical stresses are thought to play a critical role in both the initiation and progression of atherosclerotic disease, as well as in the failure of vascular reconstructions^2,15,19^. As a result, implanted stents and grafts must be designed to withstand not only hemodynamic loads but also repeated mechanical deformation over time^15^.

In recent years, efforts have been made to quantify these arterial deformations in human patients^3,8,16^ and cadaver models^4,5,17,22^. These studies provide valuable insights into the boundary conditions under which vascular repair devices must operate, guiding the design and testing of stents and other implants^20,30^. However, while cadaver models offer detailed anatomical insights, they lack the dynamic biomechanical environment and biological responses of living systems. As such, they cannot capture the full spectrum of device-tissue interactions, particularly those related to remodeling and healing. To evaluate the real-world performance, safety, and long-term durability of vascular devices, *in vivo* testing in live animal models is essential.

Large animals are preferred for preclinical testing, as they can accommodate implants designed for human anatomical dimensions. Among these, swine are widely regarded as the most suitable species due to their close anatomical and physiological similarities to humans^28,31^. Their arterial size permits the use of human-scale devices, and their cardiovascular responses closely mirror those of human patients under both baseline and pathological conditions^12,29^. Swine models have been extensively used in cardiovascular research, including for peripheral stent implantation studies^23^. However, despite their widespread application, the biomechanical compatibility of porcine and human lower extremity arteries remains poorly characterized. In particular, it is unclear whether porcine arteries experience similar magnitudes and patterns of flexion-induced compression, bending, and tortuosity, or whether these parameters vary by breed, age, or weight.

In this study, we sought to address this gap by characterizing the geometry and mechanical behavior of lower extremity arteries in two commonly used swine breeds: Yucatan minipigs, which are favored for their limited growth potential in long-term studies, and domestic swine (Sus scrofa domesticus), which are more widely available and frequently used for short-term cardiovascular research. Our objectives were to quantify arterial diameters, lengths, tortuosity, and bending characteristics in the straight and flexed limb positions, and to evaluate how these parameters are influenced by age and weight of the animals. Understanding these biomechanical features is critical for refining preclinical models of PAD, designing more robust vascular devices, and ultimately improving outcomes in patients undergoing lower extremity revascularization.

## MATERIALS AND METHODS

### Animals

All procedures were approved by the Institutional Animal Care and Use Committee (IACUC) of the University of Nebraska Medical Center and conducted in accordance with the U.S. Public Health Service Policy on Humane Care and Use of Laboratory Animals and the National Research Council’s Guide for the Care and Use of Laboratory Animals. A total of 20 Yucatan swine (Sus scrofa) (mean age: 15.9 ± 2.7 months; range: 10.6 - 22.0 months; 58 CTAs) and 16 domestic swine (Sus scrofa domesticus) (mean age: 4.5 ± 1.5 months; range: 3.0 - 8.0 months; 21 CTAs) were included in this study. The average weight of Yucatan animals was 66.7 ± 10.4 kg (range: 50.0 - 100.0 kg), while domestic swine averaged 49.2 ± 18.4 kg (range: 31.5–117.0 kg). All but one Yucatan swine underwent three CTAs, with the first and last spaced an average of 4.2 ± 0.6 months apart (range: 2.3 - 5.1 months), whereas only five domestic swine underwent two CTAs, with an average interval of 1.9 ± 1.1 months (range: 0.9 - 3.6 months). In addition, 19 Yucatan and 11 domestic swine underwent both straight and bent-leg scans. None of the animals had any interventions performed on the abdominal aorta or lower extremity arteries.

For imaging procedures, animals were sedated via intramuscular injection of Tiletamine/Zolazepam (4.4 mg/kg) combined with Ketamine (2.2 mg/kg) as a single dose. Following sedation, animals were intubated and maintained under general anesthesia using 1-5% Isoflurane delivered continuously through an endotracheal tube with a calibrated vaporizer during transport and throughout the imaging session. At the conclusion of terminal procedures, euthanasia was performed using one of two protocols: (1) intravenous administration of Pentobarbital (86.7 mg/kg) combined with Phenytoin (11.1 mg/kg) as a single dose, or (2) intra-aortic infusion of a cardioplegic solution composed of 26 mEq/L potassium chloride, 13 mL of 1% lidocaine, 3.2 g/L of 20% mannitol, 2 g of 50% magnesium sulfate, 13 mEq/L of sodium bicarbonate, and 1000 mL of Plasma-Lyte A. In all cases, euthanasia was confirmed by bilateral thoracotomy.

### Imaging and image analysis

Computed tomography angiography (CTA) scans were acquired with the hind limbs positioned either straight or bent (Figure 1). For the straight limb posture, mimicking quadrupedal stance, limbs were extended and secured with elastic tape. For the bent posture - mimicking limb positioning during sternal recumbency - the limbs were flexed and pressed against the body. Some scans captured one limb in each posture, while others included both limbs in both positions. All imaging was performed using a 256-slice GE Revolution CT scanner (GE Healthcare, Chicago, IL), yielding 512 × 512 pixel images at a resolution of 0.23 mm and an axial slice thickness of 0.625 mm. Scans were performed at 120 kVp following intravenous injection of 100 mL of iopamidol contrast (Bracco Imaging, Milan, Italy). DICOM images were imported into Materialise Mimics v25 (Materialise, Leuven, Belgium) for 3D reconstruction using standard tools for masking, region growing, mask separation, and model generation, as described previously^9,11,18^.

**Figure 1.**
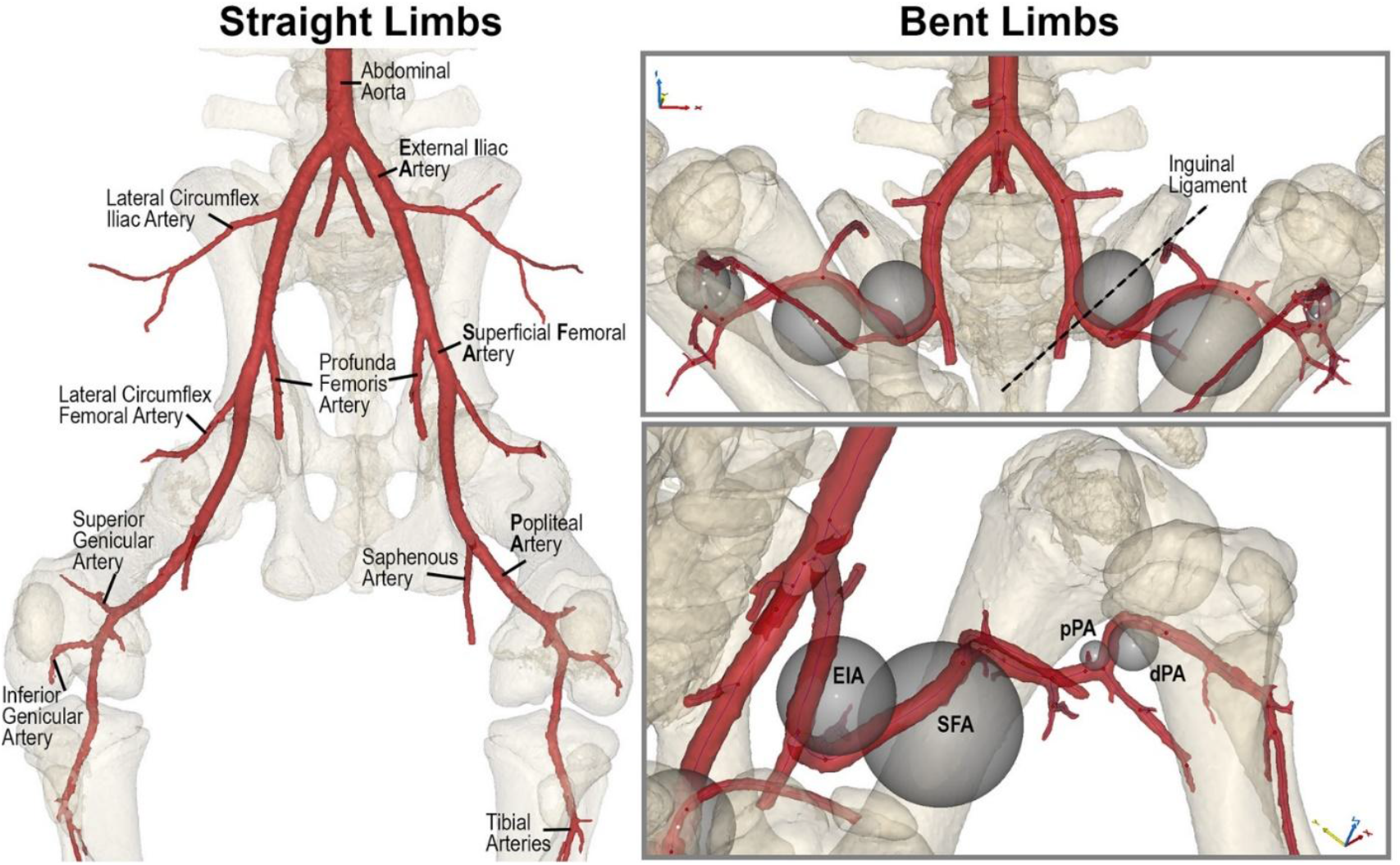
Porcine lower extremity arteries in the straight (left) and bent (right) limb postures. Inscribed sphere diameters in the bent limb position were used to characterize arterial bending. EIA = External Iliac Artery, SFA = Superficial Femoral Artery, pPA = proximal Popliteal Artery, dPA = distal Popliteal Artery.

Separate masks were created for the arterial tree and skeletal structures. The bone mask was used to measure hip and knee flexion angles in the lateral or medial planes. The arterial mask was used to extract vessel centerlines for accurate measurement of diameters and lengths, minimizing errors due to oblique slicing, as outlined in established protocols^11,14^. Arterial diameters were measured at the following standardized locations (Figure 1):

- Abdominal aorta (AA): ∼1 cm proximal to the iliac trifurcation
- Proximal external iliac artery (pEIA): ∼1 cm distal to the aortoiliac bifurcation
- Distal external iliac artery (dEIA): ∼1 cm proximal to the profunda femoris artery
- Proximal superficial femoral artery (pSFA): ∼1 cm distal to the profunda femoris artery
- Proximal popliteal artery (pPA): within 1 cm distal to the saphenous artery origin
- Distal popliteal artery (dPA): at the point of contact with the tibia, just before the artery passes through the interosseous membrane and bifurcates into the tibial arteries.

Arterial lengths in both postures were measured along the centerline between:

- the aortoiliac bifurcation and the origin of the profunda femoris artery (EIA)
- the origin of the profunda femoris and the saphenous arteries (SFA)
- the saphenous artery and the tibial arteries (PA).

Axial compression was calculated as the percentage reduction in arterial length during flexion, using the formula:

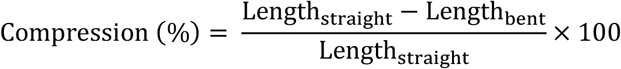

as previously described in human studies^19,22^.

Arterial tortuosity was quantified as the difference between the centerline distance and the shortest distance between the two points, divided by the centerline distance, and measured for the EIA, SFA, and PA. To assess arterial bending, we inscribed spheres within each bend (Figure 1), and the radius of each sphere was used as a quantitative measure of bend severity, following methods established in prior work^19,22^. In addition, arterial bending angles were measured in the lateral plane for the following segments: behind the inguinal ligament (EIA), in the SFA, the pPA, and the dPA.

### Statistical analysis

Results are reported as mean ± standard deviation, median, 25th and 75th percentiles, and minimum and maximum values. To assess changes in arterial diameters and lengths in relation to age and weight of the Yucatan swine, linear models were fitted for each animal across measured arterial locations, with the slope representing the rate of vascular growth for each segment. The distribution of slopes was tested for normality using the Shapiro-Wilk test. For normally distributed variables, two-tailed t-tests were applied; for non-normally distributed variables (p < 0.05), the Wilcoxon signed-rank test was used.

In domestic swine, only 5 of 16 animals underwent repeat CTA, yielding two timepoints per animal. Therefore, linear mixed-effects models were used to assess changes in arterial diameters and lengths while accounting for repeated measures. Diameters and lengths were specified as dependent variables, with age and weight as continuous fixed-effect covariates. A random intercept for Animal ID was included to account for intra-animal correlation, with no additional repeated effects structure specified. Models were fitted using restricted maximum likelihood (REML) estimation, and statistical significance was evaluated using Type III F-tests. Marginal and conditional R^2^ values were calculated to assess model performance, and 95% confidence intervals were reported for all parameter estimates.

To evaluate the effects of age, weight, hip flexion, and knee flexion on arterial biomechanics - including axial compression, tortuosity, inscribed sphere radii, and bending angles - linear mixed-effects models were similarly used. Each arterial segment and outcome variable was modeled independently. Fixed effects included standardized (z-scored) values of age, weight, hip angle, and knee angle, while random intercepts for Animal ID captured subject-specific variability. Models were again fitted using REML, and model performance was quantified using marginal R^2^ values. Conditional R^2^ values were not computed, as between-subject variance was minimal in most models.

All analyses were performed using IBM SPSS Statistics v29 (IBM Corp., Armonk, NY) with a diagonal covariance structure and variance components estimation for random effects. A p-value of less than 0.05 was considered statistically significant.

## RESULTS

### Arterial diameters

In both Yucatán and domestic swine, arterial diameters progressively decreased from the AA to the pEIA, dEIA, pSFA, pPA, and dPA (Figure 2). In Yucatan swine at baseline (13.8 ± 2.1 months; 58.3 ± 4.9 kg), arterial diameters measured 10.2 ± 0.9 mm in the AA, 6.2 ± 0.8 mm in the pEIA, 5.7 ± 0.9 mm in the dEIA, 4.6 ± 0.8 mm in the pSFA, 4.0 ± 0.6 mm in the pPA, and 3.4 ± 0.4 mm in the dPA. No significant changes in arterial diameter were observed with either age (p = 0.252 - 0.873) or weight (p = 0.334 - 0.948) across any of the measured segments.

**Figure 2.**
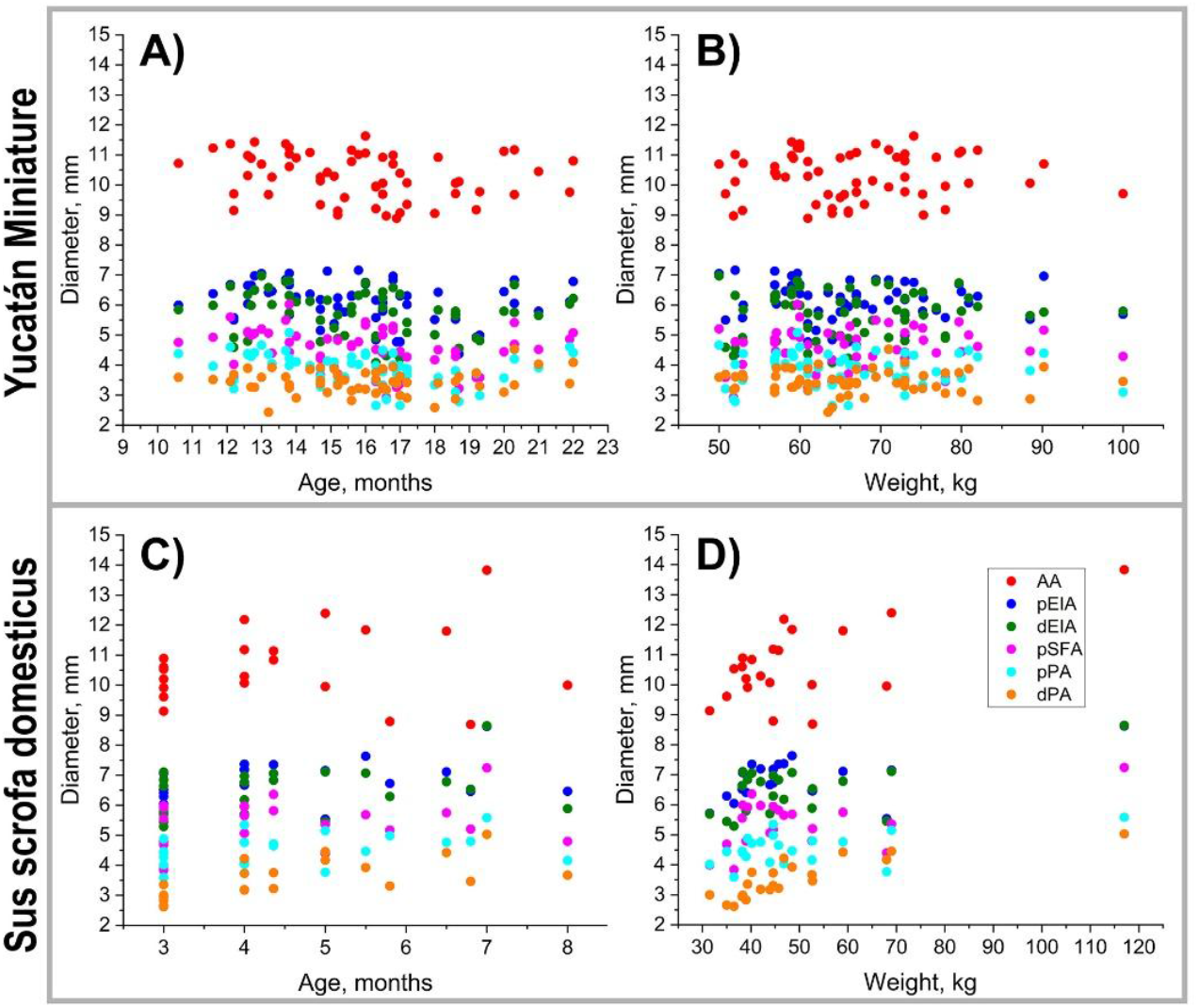
Peripheral artery diameters in Yucatán (**A, B**) and domestic (**C, D**) swine as a function of age (**A, C**) and weight (**B, D**). AA = abdominal aorta, pEIA = proximal external iliac artery, dEIA = distal external iliac artery, pSFA = proximal superficial femoral artery, pPA = proximal popliteal artery, dPA = distal popliteal artery. Yucatan: 20 animals, 58 measurements; Domestic: 16 animals, 21 measurements.

In domestic swine at baseline (3.9 ± 0.9 months; 42.6 ± 8.2 kg), arterial diameters measured 10.4 ± 0.9 mm in the AA, 6.7 ± 0.6 mm in the pEIA, 6.3 ± 0.7 mm in the dEIA, 5.3 ± 0.8 mm in the pSFA, and 3.3 ± 0.5 mm in the dPA. Age had no significant effect on arterial diameters at any location (p = 0.125 - 0.848). However, weight was positively associated with diameter increases in the AA, pEIA, dEIA, and dPA, with respective growth rates of 0.059 mm/kg (p = 0.001), 0.021 mm/kg (p = 0.048), 0.028 mm/kg (p = 0.016), and 0.024 mm/kg (p < 0.001). Correlations between weight and diameters of the pSFA and pPA were not statistically significant (p = 0.065 and 0.115, respectively).

Summary measurements of arterial diameters for each segment in both breeds at baseline, along with their corresponding growth rates (where applicable), are presented in Table 1.

**Table 1.**
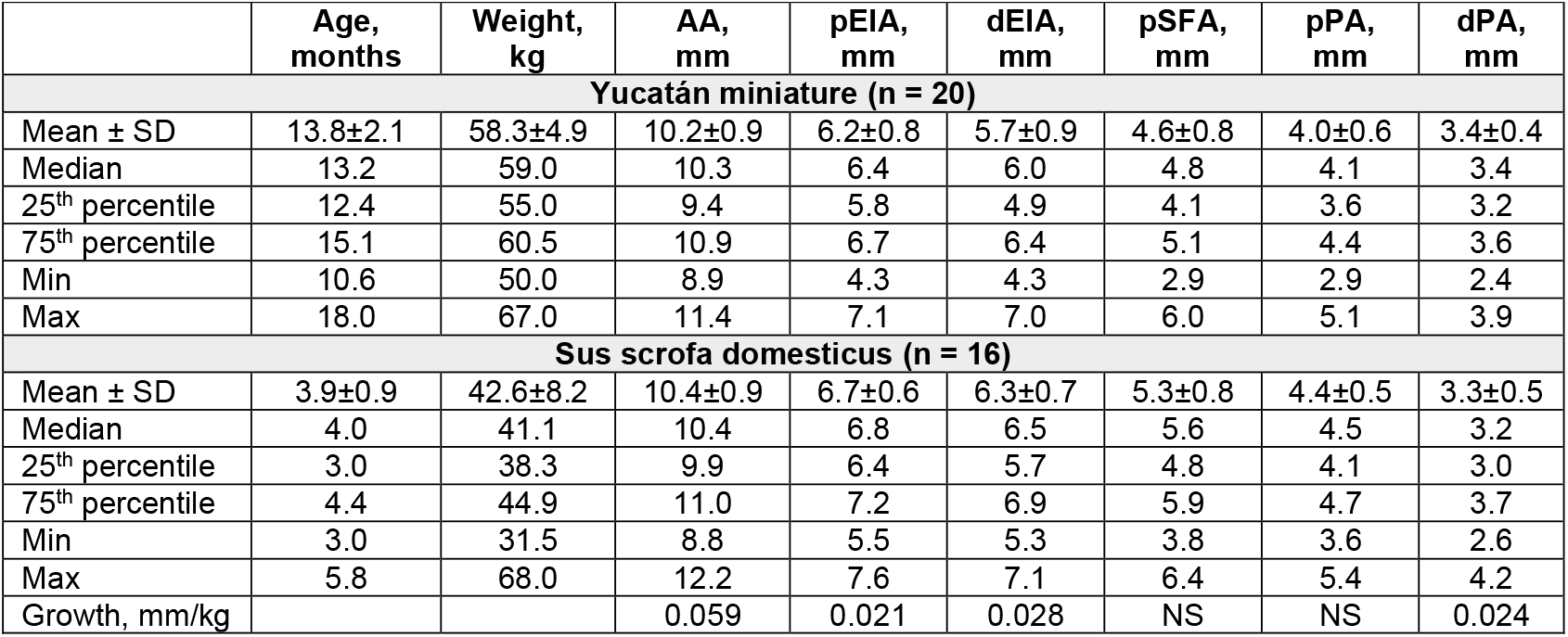
Peripheral artery diameters (mm) for Yucatan miniature and *sus scrofa domesticus* swine at baseline, along with their growth characteristics when applicable. NS = not significant; n represents the number of animals used to obtain these parameters.

### Lengths

Lengths of the EIA, SFA, and PA as functions of age and weight for both swine breeds are presented in Figure 3, with baseline measurements and growth rates summarized in Table 2. In Yucatan swine at baseline (14.0 ± 2.2 months; 59.8 ± 9.0 kg), arterial lengths in the straight limb posture measured 87 ± 9 mm in the EIA, 103 ± 8 mm in the SFA, and 123 ± 9 mm in the PA. Lengths increased significantly with age across all segments: 1.38 ± 1.61 mm/month in the EIA (p = 0.003), 1.54 ± 2.09 mm/month in the SFA (p = 0.001), and 1.46 ± 1.62 mm/month in the PA (p = 0.001). Weight-related increases were also observed in the EIA (0.44 ± 0.82 mm/kg, p = 0.003) and SFA (0.30 ± 0.42 mm/kg, p = 0.007), but not in the PA (p = 0.123).

**Table 2.** Peripheral artery lengths (mm) for Yucatan miniature and *sus scrofa domesticus* swine in the straight limb position at baseline, along with their growth characteristics when applicable. NS = not significant; n represents the number of animals used to obtain these parameters.

|  | Age, months | Weight, kg | EIA, mm | SFA, mm | PA, mm |
| --- | --- | --- | --- | --- | --- |
| **Yucatán miniature (n = 19)** |  |  |  |  |  |
| Mean ± SD, mm | 14.0±2.2 | 59.8±9.0 | 87±9 | 103±8 | 123±9 |
| Median, mm | 13.2 | 59.0 | 86 | 102 | 121 |
| 25<sup>th</sup> percentile, mm | 12.3 | 54.0 | 80 | 99 | 119 |
| 75<sup>th</sup> percentile, mm | 16.3 | 60.8 | 93 | 108 | 126 |
| Min, mm | 10.6 | 50.0 | 68 | 84 | 110 |
| Max, mm | 18.0 | 90.2 | 105 | 119 | 151 |
| Growth, mm/month | | | 1.38±1.61 | 1.54±2.09 | 1.46±1.62 |
| Growth, mm/kg | | | 0.44±0.82 | 0.30±0.42 | NS |
| **Sus scrofa domestica (n = 11)** |  |  |  |  |  |
| Mean ± SD, mm | 3.8±1.1 | 42.3±6.4 | 81±5 | 93±8 | 124±7 |
| Median, mm | 4.0 | 40.2 | 80 | 91 | 123 |
| 25<sup>th</sup> percentile, mm | 3.0 | 38.7 | 79 | 86 | 121 |
| 75<sup>th</sup> percentile, mm | 4.2 | 44.3 | 83 | 98 | 127 |
| Min, mm | 3.0 | 35.0 | 71 | 81 | 113 |
| Max, mm | 6.5 | 59.0 | 88 | 106 | 137 |
| Growth, mm/month |  |  | NS | 3.96 | 3.13 |
| Growth, mm/kg |  |  | NS | 1.00 | 0.82 |

**Figure 3.**
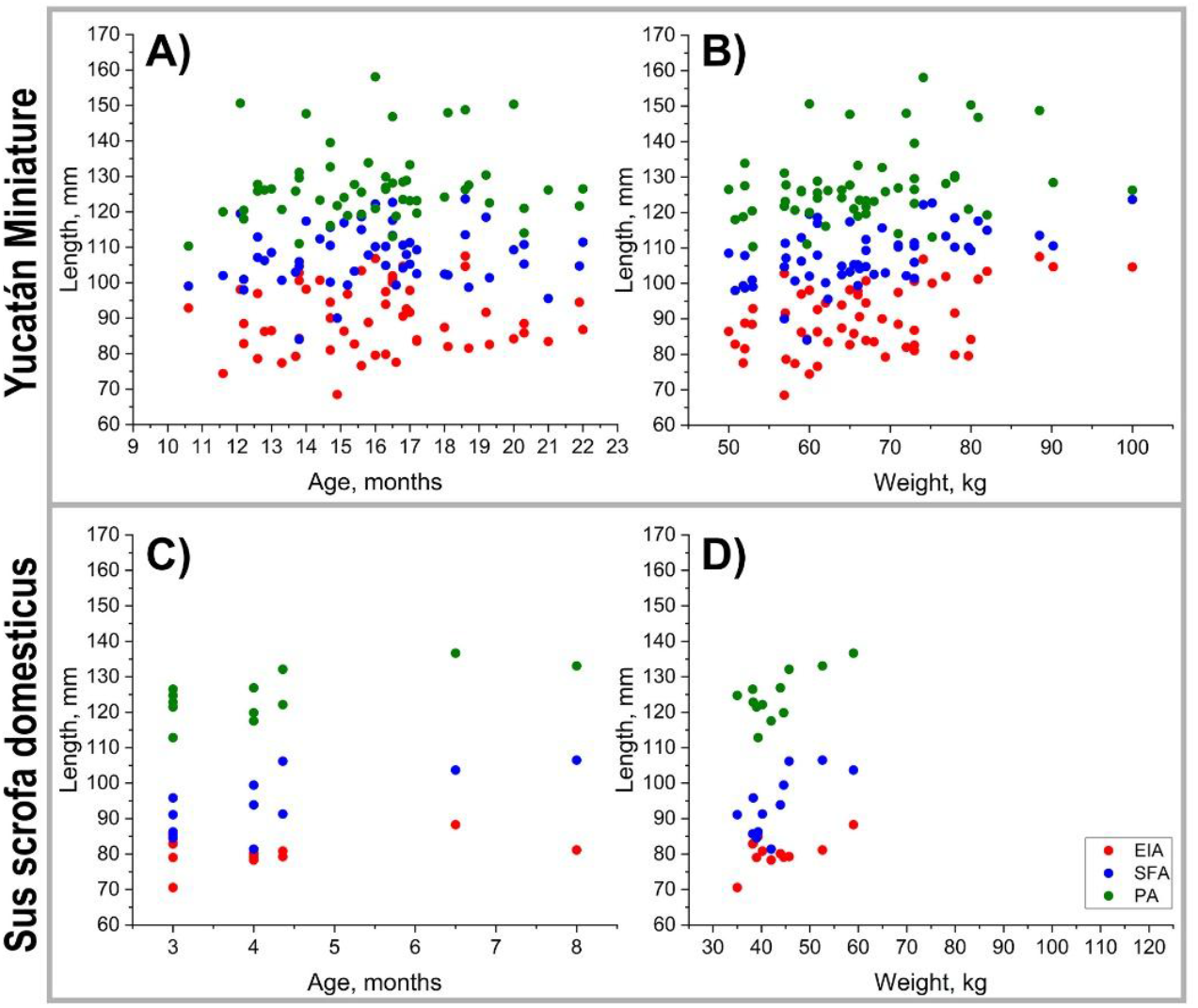
Peripheral artery lengths in Yucatán (**A, B**) and domestic (**C, D**) swine as a function of age (**A, C**) and weight (**B, D**). EIA = external iliac artery, SFA = superficial femoral artery, PA = popliteal artery. Yucatan: 19 animals, 55 measurements; Domestic: 11 animals, 12 measurements.

In domestic swine at baseline (3.8 ± 1.1 months; 42.3 ± 6.4 kg), arterial lengths in the straight limb posture measured 81 ± 5 mm in the EIA, 93 ± 8 mm in the SFA, and 124 ± 7 mm in the PA. EIA length did not change significantly with either age (p = 0.530) or weight (p = 0.214). However, lengths of the SFA and PA increased significantly with both age (p = 0.017 and p = 0.016, respectively) and weight (p = 0.011 and p = 0.007, respectively), at rates of 3.96 mm/month and 3.13 mm/month, and 1.00 mm/kg and 0.82 mm/kg, respectively.

### Axial compression and tortuosity

In Yucatan animals, limb flexion resulted in an average hip angle of 55 ± 13° (range: 42-133°, 25^th^ percentile: 48°, 75^th^ percentile: 58°) and knee angle of 47 ± 12° (range: 33-102°, 25^th^ percentile: 41°, 75^th^ percentile: 49°). In domestic swine, average hip and knee flexion angles were 52 ± 8° (range: 30 - 66°, 25^th^ percentile: 52°, 75^th^ percentile: 55°) and 52 ± 6° (range: 47 - 64°, 25^th^ percentile: 48°, 75^th^ percentile: 54°), respectively.

Compared to the straight limb position, flexion in Yucatan swine resulted in axial compression of 15.4 ± 6.4% in the EIA, 32.7 ± 4.7% in the SFA, and 14.9 ± 6.1% in the PA (Figure 4A). Compression of the SFA was significantly greater than that of both the EIA and the PA (p < 0.001), while no significant difference was observed between EIA and PA compression (p = 0.633). No significant changes in axial compression were observed with increasing age or weight in any arterial segment, except in the PA, where compression decreased with weight at a rate of -0.440 ± 0.778 %/kg (p = 0.035).

**Figure 4.**
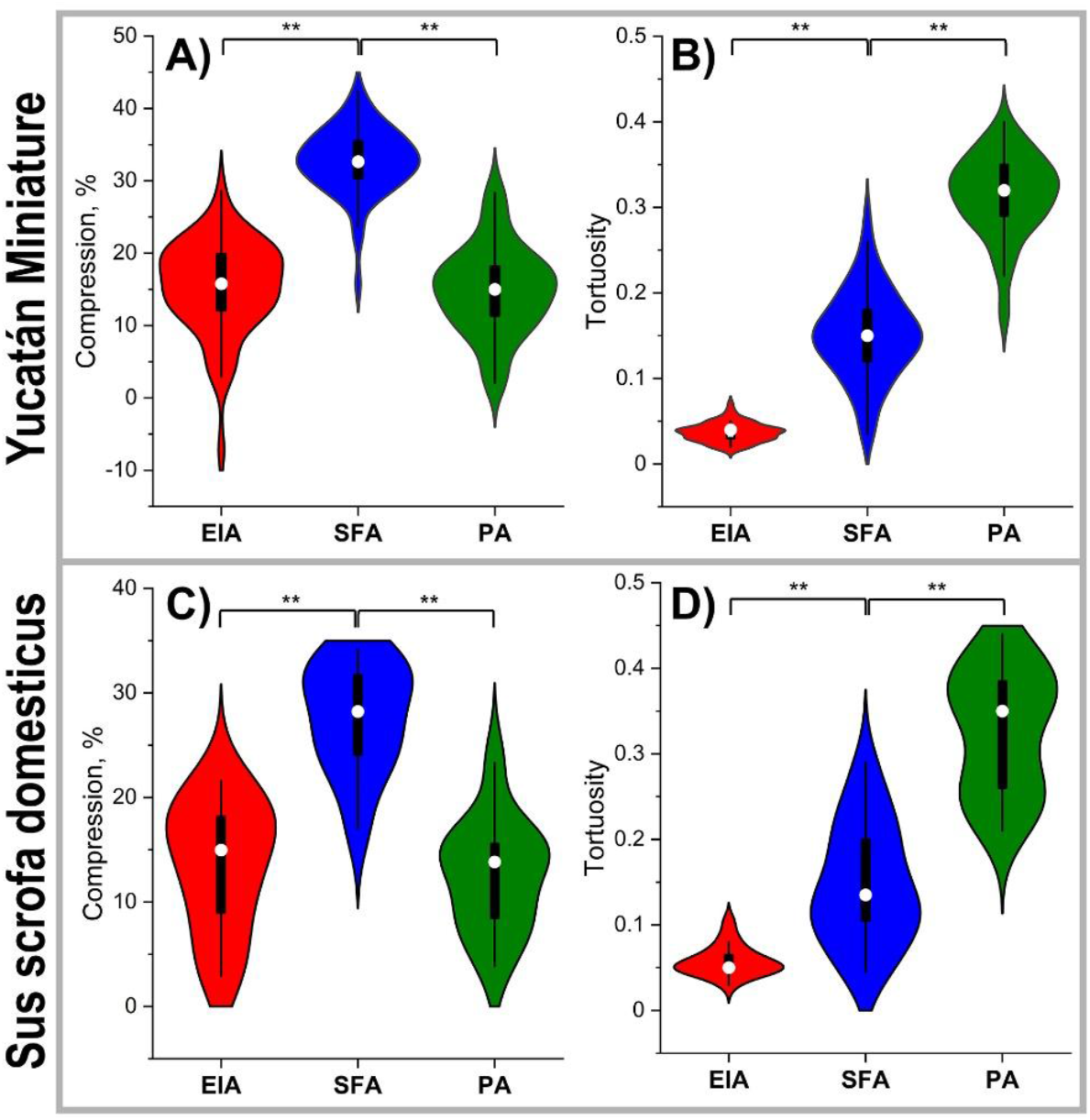
Compression (**A**,**C**) and tortuosity (**B, D**) in the external iliac artery (EIA), superficial femoral artery (SFA), and popliteal (PA) artery in the bent limb position of Yucatán (**A, B**) and domestic (**C, D**) swine. In the violin plots, the box extends to 25th and 75th percentiles, whiskers mark 1.5 standard deviations, and median values are marked with a hollow circle. Significance at p < 0.001 is marked with **. Compression: Yucatan - 19 animals, 54 measurements; Domestic: 11 animals, 12 measurements. Tortuosity: Yucatan – 20 animals, 58 measurements; Domestic – 16 animals, 21 measurements. Statistics: paired t-tests and Wilcoxon signed-rank test.

In domestic swine, axial compression was slightly lower, measuring 13.5 ± 6.1% in the EIA, 27.6 ± 5.2% in the SFA, and 12.6 ± 5.3% in the PA (Figure 4C). Similar to the findings in Yucatan swine, SFA compression in domestic animals was significantly greater than that observed in the EIA and PA (p < 0.001), while EIA and PA compression did not differ significantly (p = 0.690).

Linear mixed-effects modeling revealed that in Yucatan swine, axial compression of the EIA was significantly associated with hip flexion angle (p = 0.021), whereas no significant effects were observed for age, weight, or knee flexion. Compression of the SFA and PA was not significantly influenced by any of the tested variables. In domestic swine, no significant effects of age, weight, hip flexion, or knee flexion were observed on axial compression of the EIA (p = 0.274-0.713), SFA (p = 0.166-0.811), or PA (p = 0.245-0.888).

Arterial tortuosity in the flexed limb posture was assessed using 58 bent-leg CTAs from Yucatan swine and 21 from domestic swine. In both breeds, tortuosity progressively increased in more distal arterial segments (p < 0.001 for all comparisons; Figure 4B, D). In Yucatan animals, average tortuosity values were 0.04 ± 0.01 for the EIA, 0.15 ± 0.05 for the SFA, and 0.31 ± 0.05 for the PA. Corresponding values in domestic swine were 0.06 ± 0.02 for the EIA, 0.15 ± 0.07 for the SFA, and 0.33 ± 0.07 for the PA.

Linear mixed-effects modeling showed that in both Yucatan and domestic swine, EIA tortuosity was not significantly influenced by age, weight, or hip and knee flexion angles. In contrast, tortuosity of the SFA and PA was significantly associated with hip flexion angle (p = 0.003 and p = 0.004, respectively) in Yucatan swine, and (p = 0.025 and p = 0.024, respectively) in domestic swine. Hip flexion angle accounted for 38% and 24% of the variance in SFA and PA tortuosity in Yucatan swine, and 44% and 47% in domestic swine, respectively.

### Inscribed sphere radii and arterial bending angles

In Yucatan swine (Figure 5A), average inscribed sphere radii were 23.8 ± 8.1 mm behind the inguinal ligament, 29.8 ± 8.0 mm in the SFA, 7.2 ± 3.0 mm in the pPA, and 9.6 ± 1.8 mm in the dPA, with significant differences across these segments (p < 0.001). Sphere radii behind the inguinal ligament were significantly influenced by age (p = 0.037), weight (p = 0.013), and hip flexion angle (p < 0.001), collectively explaining 55% of the observed variance. In the SFA, sphere radii were affected by weight (p = 0.028) and hip flexion angle (p = 0.008), accounting for 40% of the variance. In the pPA, only hip flexion angle had a significant effect (p < 0.001), explaining 16% of the variance. For the dPA, knee flexion angle was the only significant predictor (p = 0.012), accounting for 34% of the variance.

**Figure 5.**
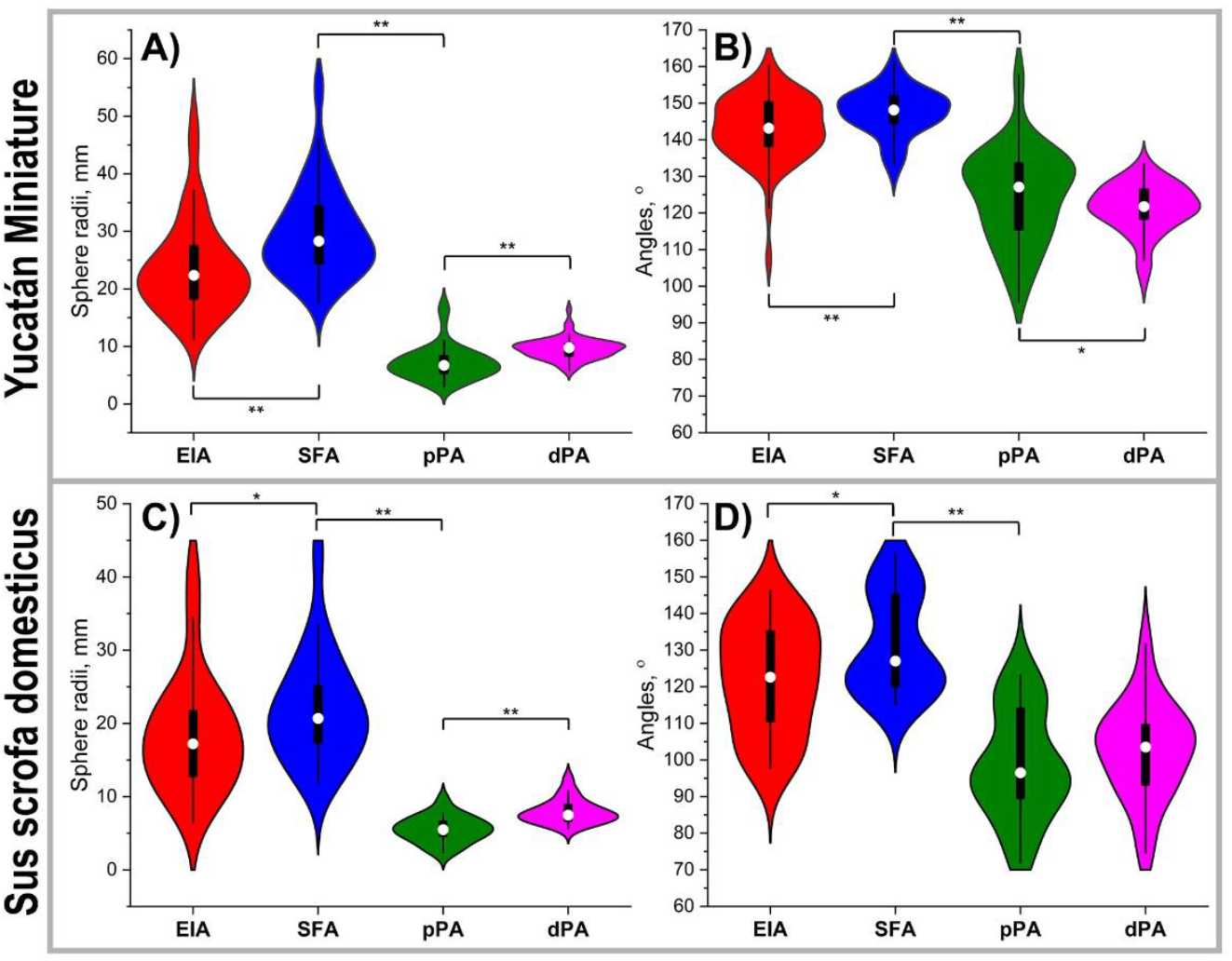
Inscribed sphere radii (**A**,**C**) and angles (**B, D**) in the external iliac artery (EIA), superficial femoral artery (SFA), and proximal (pPA) and distal (dPA) popliteal artery in the bent limb position of Yucatán (**A, B**) and domestic (**C, D**) swine. In the violin plots, the box extends to 25th and 75th percentiles, whiskers mark 1.5 standard deviations, and median values are marked with a hollow circle. Significance at p < 0.001 is marked with **, and at p < 0.05 with *. Yucatan: 20 animals, 58 measurements; Domestic: 16 animals, 21 measurements. Statistics: paired t-tests and Wilcoxon signed-rank test.

In domestic swine (Figure 5C), the corresponding sphere radii were slightly smaller: 18.5 ± 8.1 mm behind the inguinal ligament, 22.0 ± 7.5 mm in the SFA, 5.6 ± 1.8 mm in the pPA, and 8.0 ± 1.7 mm in the dPA. Segmental differences were also statistically significant in this breed (p = 0.024 and p < 0.001). Sphere radii behind the inguinal ligament were significantly influenced by weight (p < 0.001) and knee flexion angle (p = 0.034), together explaining 66% of the variance. In the SFA, animal weight was the only significant predictor (p = 0.006), explaining 76% of the variance in sphere radii. In the pPA and dPA, none of the tested variables - age, weight, hip flexion, or knee flexion - had a significant effect on sphere radii.

Average arterial bending angles in Yucatan swine (Figure 5B) were 143 ± 9° behind the inguinal ligament, 148 ± 6° in the SFA, 125 ± 13° in the pPA, and 121 ± 6.3° in the dPA. Differences across segments were statistically significant (p < 0.001), including between the pPA and dPA (p = 0.025). Bending angles behind the inguinal ligament, in the SFA, and in the dPA were not significantly influenced by age, weight, or hip and knee flexion angles. In contrast, the bending angle at the pPA was significantly affected by both weight (p = 0.031) and hip flexion angle (p = 0.013), which together explained 52% of the observed variance.

In domestic swine (Figure 5D), arterial bending angles were consistently smaller across all segments compared to Yucatan swine (p < 0.001 for all locations), with average values of 122 ± 15° behind the inguinal ligament, 132 ± 14° in the SFA, 99 ± 15° in the pPA, and 103 ± 13° in the dPA. As in Yucatans, segmental differences in arterial angle were statistically significant (p < 0.001), except between the pPA and dPA, which did not differ significantly (p = 0.146). In this breed, bending angles behind the inguinal ligament were significantly influenced by both age (p = 0.035) and weight (p = 0.029), explaining 37% of the variance. Bending angles in the pPA and dPA were significantly associated with hip flexion (p = 0.046) and knee flexion (p = 0.037), explaining 36% and 34% of the variance, respectively.

## DISCUSSION

PAD remains a significant clinical challenge, particularly in the lower extremities, where the combination of lesion location and complex biomechanical deformations during limb flexion complicates both disease progression and treatment outcomes. The development and optimization of vascular repair strategies - including stents, bypass grafts, drug-coated balloons, plaque modification techniques such as atherectomy or intravascular lithotripsy, and bioresorbable scaffolds - require robust preclinical models that faithfully replicate both the anatomical dimensions and the biomechanical environment of human lower limb arteries. In this study, we systematically characterized the geometry and biomechanical behavior of the EIA, SFA, and PA in two commonly used swine breeds. Our results demonstrate that swine closely replicate key features of human lower extremity arterial biomechanics, including vessel diameters, segment lengths, flexion-induced compression, and bending, making them a highly relevant large-animal model for PAD research and device evaluation.

Specifically, our analysis indicates that EIA diameters in Yucatan swine at 12 to 15 months of age were 4.9 – 6.7 mm, while those in domestic swine were slightly larger, ranging from 5.7 to 7.2 mm even at 4 months of age, and continued to grow. These values closely correspond to human SFA diameters^25^, which typically range from 5 to 7.5 mm. The porcine SFA measured 4.1-5.1 mm in Yucatan swine and 4.8-5.9 mm in domestic swine, aligning more closely with the dimensions of the human PA^10^ (4.0-6.5 mm). In contrast, the porcine PA measured 3.2-4.4 mm in Yucatans and 3.0-4.7 mm in domestic swine, which is more consistent with the size of human tibial arteries^27^, typically ranging from 2.0 to 3.5 mm.

The lengths of the porcine arterial segments ranged from 79-93 mm for the EIA, 86-108 mm for the SFA, and 119-127 mm for the PA in animals aged 12-16 months (Yucatan) and 3-4 months (domestic swine). These lengths continued to increase with age and weight. Notably, these dimensions are comparable to those of the human PA and are more than sufficient to support the evaluation of vascular repair devices, including configurations involving multiple overlapping stents. This is particularly important because, in humans, the femoropopliteal artery segment that undergoes the most severe mechanical deformations - and is most commonly affected by atherosclerotic disease^32^ - is located around the adductor hiatus and extends into the PA behind the knee^17,19,22^. The porcine arterial segments analyzed in this study closely match the length and anatomical context of these high-risk human regions, further supporting their utility in preclinical testing of vascular and endovascular devices.

Importantly, both swine breeds replicated the worst-case scenario of human limb flexion - namely, the fetal posture characterized by acutely bent limbs - exhibiting knee flexion angles of 47-52°, which closely approximate the 50-60° observed in humans under similar conditions. In the flexed posture, porcine arteries exhibited 12-33% axial compression, with comparable values observed across both swine breeds. The highest compression occurred in the SFA, making it a representative worst-case scenario for axial deformation typically seen in humans. Specifically, human studies have reported up to 38% compression in the PA during fetal posture, with the 25^th^ and 75^th^ percentiles ranging from 21% to 29%^22^. In comparison, the porcine SFA undergoes 25-35% compression under similar conditions, with maximum values reaching 42%. Additionally, the porcine EIA and PA exhibit deformation patterns analogous (13-15%) to those observed in the human SFA and adductor hiatus regions during sitting and fetal postures, where average compression in humans ranges from 13% to 19%^22^. These findings are also consistent with measurements in PAD patients, in whom the femoropopliteal artery exhibits an average of 9 ± 3% axial compression in the sitting posture^13^.

Similarly to human lower extremity arteries, tortuosity in swine increased with more distal arterial locations, reaching 0.31-0.33 in the PA compared to 0.15 in the SFA and 0.04-0.06 in the EIA. Arterial bending radii were smallest in the PA, ranging from 6 to 10 mm, with slightly more acute bending observed in the proximal segment compared to the distal segment. These values closely align with human data^22^, where bending radii of 8 - 11 mm have been reported in the fetal and sitting postures, with minimum radii reaching as low as 5 mm. In the porcine SFA and EIA, particularly behind the inguinal ligament, bending radii ranged from 19 to 30 mm. These measurements are consistent with values reported in human cadaver studies of the SFA and adductor hiatus during walking and sitting postures (12–27 mm), and also correspond well with those observed in PAD patients^13^. Bending angles in swine mirrored the pattern observed in sphere radii, with the most acute angulations occurring in the PA, ranging from 99° to 125°. These values closely correspond to those reported in human PAs during the fetal posture, which exhibit an average bending angle of 136° and a minimum of 117° ± 17°. The porcine PA bending angles also align well with measurements in PAD patients in the sitting posture, where mean angles of 101° ± 22° have been documented^13^.

An important consideration when selecting an appropriate animal model for preclinical testing is the breed’s growth potential, typically evaluated in terms of age and body weight. In Yucatan miniature swine, vessel diameters remained stable across analyzed age and weight, although both were associated with increased arterial lengths. In contrast, domestic swine showed a significant relationship between body weight and both vessel diameters and lengths. These findings highlight the importance of breed selection - particularly for studies requiring consistent vascular dimensions - and suggest that Yucatan swine may be preferable for long-term or serial studies due to their limited growth potential. Interestingly, in both breeds, axial compression during limb flexion and arterial tortuosity were not significantly affected by age or weight. However, inscribed sphere radii - used to characterize vessel bending - did show some dependence on these variables in more proximal segments, likely reflecting generalized somatic growth and corresponding increases in vessel size, rather than localized biomechanical changes.

While this study provides a comprehensive analysis of the lower extremity vasculature in two commonly used swine breeds, several limitations should be considered when interpreting the results. First, the number of domestic swine included in the study was smaller than that of Yucatan swine (16 vs. 20 animals), and domestic animals underwent fewer repeat scans. As a result, a different analytical approach was required to assess growth in this group, limiting the direct comparability of growth trends across breeds. Additionally, a substantial gap in body weight was observed among domestic swine between 70 and 117 kg, which may confound the generalizability of weight-associated growth patterns in this group. Nonetheless, data from animals weighing 30-70 kg still demonstrated appreciable growth trends. Second, although the animals included in this analysis covered a representative range of ages and weights - 11 to 22 months for Yucatans and 3 to 8 months for domestic swine - this range, while typical for preclinical research, does not encompass the full spectrum of growth and maturation, particularly in rapidly growing domestic breeds. Future studies should consider expanding the scope to include additional breeds, such as Ossabaw swine, which offer unique metabolic and anatomical profiles relevant to vascular disease modeling^12^. Furthermore, this study focused exclusively on morphological and biomechanical parameters derived from imaging. Follow-up investigations incorporating biochemical characterization, such as lipid profiles or inflammatory markers, as well as direct measurements of arterial mechanical properties, would provide a more holistic understanding of the vascular phenotype and its relevance to human PAD.

## DECLARATIONS

### Ethics approval and consent to participate

All procedures were approved by the Institutional Animal Care and Use Committee (IACUC) of the University of Nebraska Medical Center and conducted in accordance with the U.S. Public Health Service Policy on Humane Care and Use of Laboratory Animals and the National Research Council’s Guide for the Care and Use of Laboratory Animals.

### Clinical trial number

Not applicable.

### Consent for publication

Not applicable.

### Availability of data and materials

The datasets used and/or analyzed during the current study are available from the corresponding author on reasonable request.

### Competing interests

The authors have no relevant disclosures.

### Funding

This work was supported in part by the NIH awards HL125736, HL147128, HL180371, and P20GM152301.

### Authors’ contributions

Conception and design: JM, AK

Acquisition of data: BO, FH, AD, KM, AK

Analysis of data: BO, AD, AK

Interpretation of data: AK, JM

Drafting and revising the manuscript: BO, FH, AD, JM, AK

Approval of the submitted version: BO, FH, AD, JM, AK

## Acknowledgements

The authors would also like to thank the Tissue Analysis Core (TAC) of the NIH Center for Cardiovascular Research in Biomechanics (CRiB) for assistance with animal procedures.

## Notes

### Competing Interest Statement

The authors have declared no competing interest.

## REFERENCES

1. Ansari, F., L. K. Pack, S. S. Brooks, and T. M. Morrison. Design considerations for studies of the biomechanical environment of the femoropopliteal arteries. J. Vasc. Surg. 58:804–813, 2013.

2. Bapat, G. M., A. Z. Bashir, P. Malcolm, J. M. Johanning, I. I. Pipinos, and S. A. Myers. A biomechanical perspective on walking in patients with peripheral artery disease. Vasc. Med. Lond. Engl. 28:77–84, 2023.

3. Cheng, C., N. Wilson, and R. Hallett. In vivo MR angiographic quantification of axial and twisting deformations of the superficial femoral artery resulting from maximum hip and knee flexion. J Vasc Interv Radiol 17:979–987, 2006.

4. Desyatova, A., W. Poulson, P. Deegan, C. Lomneth, A. Seas, K. Maleckis, J. MacTaggart, and A. Kamenskiy. Limb flexion-induced twist and associated intramural stresses in the human femoropopliteal artery. J. R. Soc. Interface 14:20170025, 2017.

5. Desyatova, A., W. Poulson, J. MacTaggart, K. Maleckis, and A. Kamenskiy. Cross-sectional pinching in human femoropopliteal arteries due to limb flexion, and stent design optimization for maximum cross-sectional opening and minimum intramural stresses. J. R. Soc. Interface 15:10–14, 2018.

6. Eid, M. A., K. Mehta, J. A. Barnes, Z. Wanken, J. A. Columbo, D. H. Stone, P. Goodney, and M. Mayo Smith. The global burden of peripheral artery disease. J. Vasc. Surg. 77:1119-1126.e1, 2023.

7. Farber, A. et al. Surgery or Endovascular Therapy for Chronic Limb-Threatening Ischemia. N. Engl. J. Med. 387:2305–2316, 2022.

8. Gökgöl, C., N. Diehm, L. Kara, and P. Büchler. Quantification of popliteal artery deformation during leg flexion in subjects with peripheral artery disease: a pilot study. J. Endovasc. Ther. Off. J. Int. Soc. Endovasc. Spec. 20:828–35, 2013.

9. Jadidi, M., W. Poulson, P. Aylward, J. MacTaggart, C. Sanderfer, B. Marmie, M. Pipinos, and A. Kamenskiy. Calcification prevalence in different vascular zones and its association with demographics, risk factors, and morphometry. Am. J. Physiol. Heart Circ. Physiol. 320:H2313–H2323, 2021.

10. Jadidi, M., S. A. Razian, E. Anttila, T. Doan, J. Adamson, M. Pipinos, and A. Kamenskiy. Comparison of morphometric, structural, mechanical, and physiologic characteristics of human superficial femoral and popliteal arteries. Acta Biomater. 121:431–443, 2021.

11. Kamenskiy, A., D. Miserlis, P. Adamson, M. Adamson, T. Knowles, J. Neme, P. Koutakis, N. Phillips, I. Pipinos, and J. MacTaggart. Patient demographics and cardiovascular risk factors differentially influence geometric remodeling of the aorta compared with the peripheral arteries. Surgery 158:1617–1627, 2015.

12. Kamenskiy, A., B. de Oliveira, F. Heinis, P. Renavikar, J. Eberth, and J. MacTaggart. Large Animal Model of Controlled Peripheral Artery Calcification. Acta Biomater. 199:301–314, 2025.

13. Kamenskiy, A., I. Pipinos, Y. Tian, B. de Oliveira, D. Orton, X. Liu, and J. MacTaggart. Vascular Calcification Amplifies Limb Flexion-Induced Arterial Deformations and Causes Intermittent Vessel Pinching. Ann. Surg. Accepted:, 2025.

14. Kamenskiy, A. V., I. I. Pipinos, J. S. Carson, J. N. Mactaggart, and B. T. Baxter. Age and disease-related geometric and structural remodeling of the carotid artery. J. Vasc. Surg. 62:, 2015.

15. Keiser, C., K. Maleckis, P. Struczewska, M. Jadidi, J. MacTaggart, and A. Kamenskiy. A method of assessing peripheral stent abrasiveness under cyclic deformations experienced during limb movement. Acta Biomater. Online ahe:, 2022.

16. Klein, A. J., S. J. Chen, J. C. Messenger, A. R. Hansgen, M. E. Plomondon, J. D. Carroll, and I. P. Casserly. Quantitative assessment of the conformational change in the femoropopliteal artery with leg movement. Catheter Cardiovasc Interv 74:787– 798, 2009.

17. MacTaggart, J. N., N. Y. Phillips, C. S. Lomneth, I. I. I. Pipinos, R. Bowen, B. Timothy Baxter, J. Johanning, G. Matthew Longo, A. S. Desyatova, M. J. Moulton, Y. A. Dzenis, and A. V. Kamenskiy. Three-dimensional bending, torsion and axial compression of the femoropopliteal artery during limb flexion. J. Biomech. 47:2249– 2256, 2014.

18. MacTaggart, J. N., W. E. Poulson, M. Akhter, A. Seas, K. Thorson, N. Y. Phillips, A. S. Desyatova, and A. V. Kamenskiy. Morphometric roadmaps to improve accurate device delivery for fluoroscopy-free resuscitative endovascular balloon occlusion of the aorta. J. Trauma Acute Care Surg. 80:941–6, 2016.

19. MacTaggart, J., W. Poulson, A. Seas, P. Deegan, C. Lomneth, A. Desyatova, K. Maleckis, and A. Kamenskiy. Stent Design Affects Femoropopliteal Artery Deformation. Ann. Surg. 270:180–187, 2019.

20. Maleckis, K., E. Anttila, P. Aylward, W. Poulson, A. Desyatova, J. MacTaggart, and A. Kamenskiy. Nitinol Stents in the Femoropopliteal Artery: A Mechanical Perspective on Material, Design, and Performance. Ann. Biomed. Eng. 46:, 2018.

21. Mustapha, J. A., S. M. Finton, L. J. Diaz-Sandoval, F. A. Saab, and L. E. Miller. Percutaneous transluminal angioplasty in patients with infrapopliteal arterial disease. Circ. Cardiovasc. Interv. 9:1–10, 2016.

22. Poulson, W., A. Kamenskiy, A. Seas, P. Deegan, C. Lomneth, and J. MacTaggart. Limb flexion-induced axial compression and bending in human femoropopliteal artery segments. J. Vasc. Surg. 67:607–613, 2018.

23. Sakaoka, A., J. Souba, S. D. Rousselle, T. Matsuda, A. Tellez, H. Hagiwara, K. Nagano, and M. Tasaki. Different Vascular Responses to a Bare Nitinol Stent in Porcine Femoral and Femoropopliteal Arteries. Toxicol. Pathol. 019262331880072, 2018.doi:10.1177/0192623318800726

24. Schillinger, M., S. Sabeti, C. Loewe, P. Dick, J. Amighi, W. Mlekusch, O. Schlager, M. Cejna, J. Lammer, and E. Minar. Balloon angioplasty versus implantation of nitinol stents in the superficial femoral artery. N. Engl. J. Med. 354:1879–88, 2006.

25. Shahbad, R., M. Pipinos, M. Jadidi, A. Desyatova, J. Gamache, J. MacTaggart, and Kamenskiy. Structural and Mechanical Properties of Human Superficial Femoral and Popliteal Arteries. Ann. Biomed. Eng., 2024.doi:10.1007/s10439-023-03435-3

26. Song, P., D. Rudan, Y. Zhu, F. J. I. Fowkes, K. Rahimi, F. G. R. Fowkes, and I. Rudan. Global, regional, and national prevalence and risk factors for peripheral artery disease in 2015: an updated systematic review and analysis. Lancet Glob. Health 7:e1020–e1030, 2019.

27. Struczewska, P., S. A. Razian, K. Townsend, M. Jadidi, R. Shahbad, E. Zamani, J. Gamache, J. MacTaggart, and A. Kamenskiy. Mechanical, Structural, and Physiologic Differences Between Above and Below-Knee Human Arteries. Acta Biomater. 177:278–299, 2024.

28. Sturek, M., M. Alloosh, and F. W. Sellke. Swine Disease Models for Optimal Vascular Engineering. Annu. Rev. Biomed. Eng. 22:25–49, 2020.

29. Suzuki, Y., A. C. Yeung, and F. Ikeno. The Representative Porcine Model for Human Cardiovascular Disease. J. Biomed. Biotechnol. 2011:195483, 2011.

30. Vogel, J., B. T. Berg, J. Dawson, and A. Kamenskiy. Flexible Implantable Device Shape History. In: Measuring the Physiologic Use Conditions of Medical Devices, edited by W. Baxter, and R. Lahm. Cham: Springer International Publishing, 2024, pp. 125–160.doi:10.1007/978-3-031-62764-4_7

31. Walters, E. M., and R. S. Prather. Advancing Swine Models for Human Health and Diseases. Mo. Med. 110:212–215, 2013.

32. Watt, J. K. J. Origin of femoro-popliteal occlusions. Br. Med. J. 2:1455–1459, 1965.

